# Optimization of AAV tools to target Müller glial cells for retinal gene therapy

**DOI:** 10.64898/2026.04.09.717359

**Authors:** Daniel Urrutia-Cabrera, Genevieve Huppert, Stephanie Chu, Luozixian Wang, Cat Binh Nguy, Chia Fei Liu, Leszek Lisowki, Chi D. Luu, Jiang-Hui Wang, Sandy Hung, Alex W. Hewitt, Chien-Ling Huang, Tom Edwards, Keith R. Martin, Raymond C.B. Wong

## Abstract

Reprogramming of Müller glial (MG) cells into retinal neurons has the potential to treat vision loss by regenerating the retina. Development of efficient gene delivery systems to target the MG cells is critical. Adeno-associated virus (AAV) serotypes and promoter specificity are important factors that influence AAV transduction profile in the retina. However, studies that optimize these parameters to specifically target MG cells are limited, in particular in rats which are commonly used for eye research. Here we tested 4 AAV serotypes and 14 promoters to optimize gene delivery to human MG cells in vitro and/or rat MG cells in vivo. We showed that the combinatorial use of MG-specific serotypes and promoters achieved high specificity for MG cell targeting, with ShH10Y serotype and the GFAP (gfaABC1D) promoter as the best performing tool to target rat MG cells in vivo. We developed new AAV vectors using known and novel MG-specific promoters and engineered short promoter variants to improve the cargo capacity of AAV delivery. Our results highlighted a number of promoters that can target MG cells in vitro or in vivo. This study further expands the AAV toolbox to target MG cells, which has important implications for retinal gene therapy development.

## Introduction

Retinal degeneration is one of the major causes of vision loss worldwide, affecting millions of people with conditions such as age-related macular degeneration (AMD), glaucoma, inherited retinal diseases (IRD) and diabetic retinopathy. Gene therapy provides a promising strategy to treat retinal diseases that are currently incurable, and adeno-associated viruses (AAVs) have emerged as one of the most widely used vectors to deliver gene therapies to the retina. Notably, Luxturna (Voretigene neparvovec) became the first AAV-based retinal gene therapy to be approved by therapeutic regulatory organizations in many countries. Luxturna is a treatment for retinal degeneration caused by *RPE65* mutation including in Leber’s congenital amaurosis and retinitis pigmentosa, which uses the AAV2 serotype to deliver a healthy copy of the gene *RPE65* through subretinal injections^1^.

Recombinant AAVs have a favourable safety profile in the retina, with minimal risk of genome integration^2^ and relatively low immunogenicity^3^. Advances in AAV engineering allowed us to target particular retinal cells with unprecedented specificity. In particular, the capsid of different AAV serotypes affects their immunogenicity, stability and transduction profiles^4^. AAV2 is the most commonly studied serotype to target the retina, since it has shown effective transduction of retinal neurons, the retinal pigment epithelium (RPE) and glial cells^5,6^, whereas other AAV serotypes exhibit a different transduction profile in the retina. Also, numerous studies have shown that the route of delivery plays a critical role in determining the AAV transduction profile^7,8^. Anatomical barriers within the eye can affect the transduction profile of AAVs, which include the vitreous, inner limiting membrane (ILM), retinal cell density and the subretinal space. To address this, new AAV capsids were engineered to improve transduction properties, such as increased transduction yield through intravitreal (IVT) injections (e.g. 7m8^9^ and GL^10^) or higher specificity for specific cell types (ShH10^11^, Anc80^12^).

In recent years, the Müller glial (MG) cells have emerged as an interesting target for regenerative therapies. Interestingly, the MG cells of some vertebrates like teleost fish can dedifferentiate into retinal progenitors upon injury and regenerate the lost retinal cells^13^. Similar regenerative ability is also observed in amphibians and chicken to some extent^14^. However, in mammals the regenerative capacity of MG cells is limited. Notably, recent cellular reprogramming studies in rodents have highlighted the potential of using reprogramming genes to stimulate MG cells to become retinal neurons. For instance, the transcription factor *Ascl1* has been extensively studied in regenerative strategies harnessing the therapeutic potential of MG reprogramming ^15,16^. The *in vivo* reprogramming of MG cells offers the remarkable possibility of regenerating retinal cells *in situ*, which allows regenerative treatments to potentially overcome some of the limitations associated with cell-based therapies that require transplantation.

To realize this potential, the development of gene delivery methods that can specifically target Müller cells has become a critical factor to ensure the efficacy and safety of these regenerative treatments. Directed evolution studies using libraries of AAV capsids have identified new AAV serotypes that have higher affinity for MG transduction, such as the AAV6 variant ShH10^11^ and the AAV2 variant M4^17^. However, these serotypes might also target other retinal cell types, albeit less efficiently compared to MG cells. The combinatorial use of MG-specific promoters and AAV serotypes can further improve the specificity of targeted gene expression in MG cells. While previous work has analyzed the transduction efficacy of AAV tools in mouse Müller cells^18,19^, serotype and promoter testing in rat models is understudied.

In this study, we analyzed the specificity for MG cell transduction for 4 AAV serotypes: AAV2, M4, ShH10^11^ and its variant ShH10Y^20^. We assessed 13 human promoters in their ability to drive gene expression in MG cells, including *GFAP*, *GLAST*, *HES1, TRDN* and *CP* and their novel engineered variants. Promoter specificity was analyzed using *in vitro* human MG model and IVT injections to the rat retina. Our results expand the tool kit of AAV systems that can be used for regenerative therapies targeting MG cells.

## Results

### Müller glia transduction using different AAV serotypes in rats

Here we investigated the transduction profile of four AAV serotypes AAV2, ShH10, ShH10Y and M4 to assess their ability to target MG cells. To increase the specificity of transgene expression to MG cells, we paired different AAV serotypes with the short glial-specific promoter GFAP (gfaABC1D) to drive GFP expression and intravitreally injected AAV into SD rats *in vivo* (Figure 1A). Immunohistochemistry analysis of GFP expression showed that all four capsids were capable of transducing MG cells, which exhibited classical radial morphology and spatial patterning (Figure 1B-E). However, GFP expression was also detected in some cells within the ganglion cell layer (GCL). To confirm the specificity of GFP expression in MG cells, we assessed the co-localization of GFP with the Müller glia maker CRALBP to determine AAV specificity (Figure 1F). Notably, our results showed that ShH10Y achieved very high specificity in MG cells (95 ± 1.6%) followed by ShH10 (91 ± 1%), which were significantly higher than a 75 ± 1.9% specificity in wild-type AAV2 serotype (Figure 1F). The AAV2 variant M4 showed an increased specificity for transducing rat MG cells compared to the parental AAV2 capsid (97 ± 2.1%), rarely targeting other retinal cells when it was paired with the *GFAP* promoter. However, the transduction efficiency for M4 using IVT injection is low, with only a small number of GFP+ MG cells detected in vivo. Also, our negative controls using sham (saline) injection and IgG isotype control were clean (Supplementary figure 1-2). These results expand on previous reports^11,17,20^ and confirmed that all four serotypes AAV2, ShH10, ShH10Y and M4 can target rat MG cells using IVT injections. Notably, the engineered capsids ShH10, ShH10Y and M4 paired with the *GFAP* promoter demonstrated significantly higher specificity for targeting Müller cells compared to AAV2.

**Figure 1:**
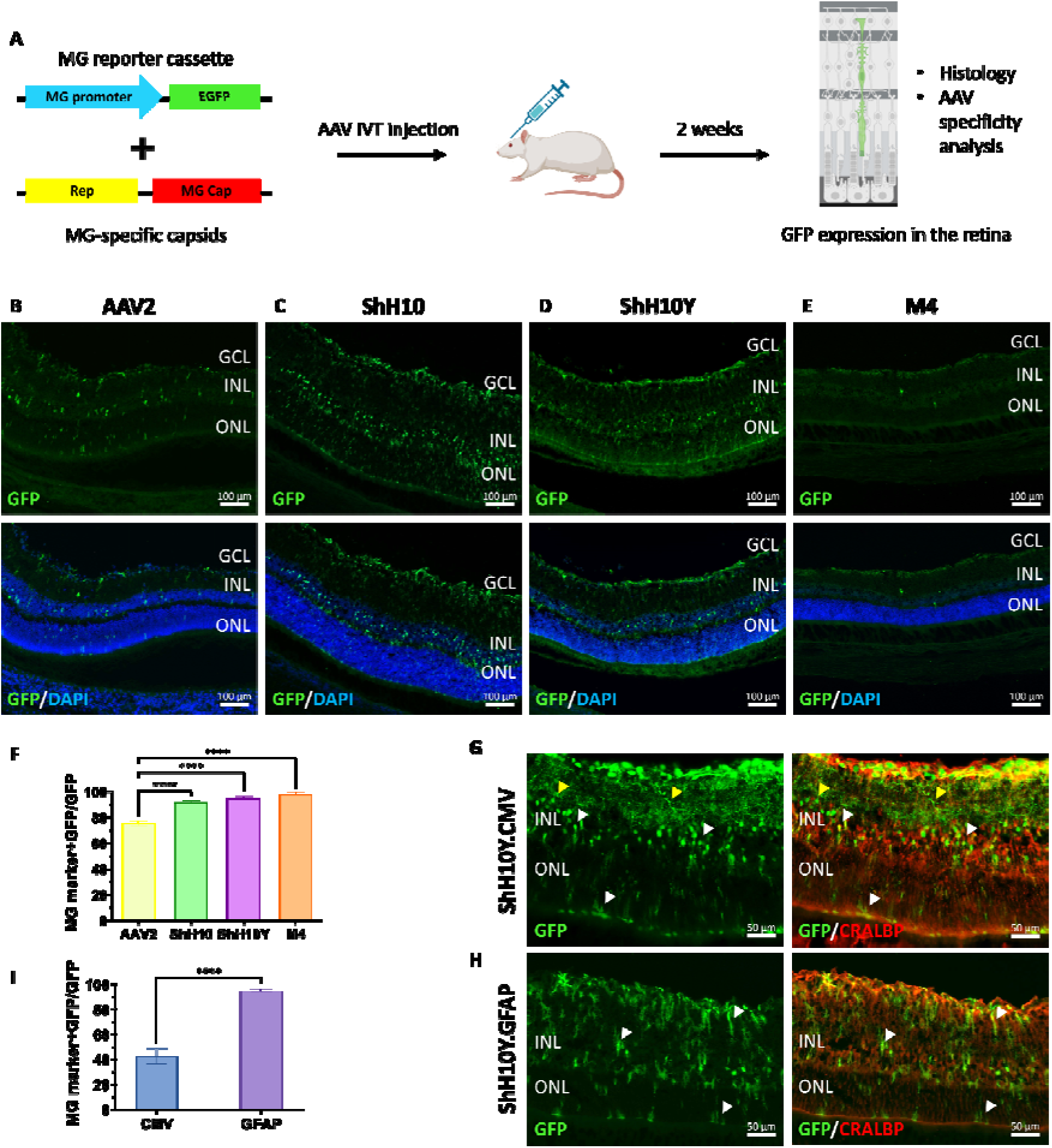
Müller glia transduction *in vivo* using different AAV capsids. A) Representative images showing the expression pattern of GFP in rat retinae transduced with IVT injections of B) AAV2.GFAP-EGFP, C) ShH10.GFAP-EGFP, D) ShH10Y.GFAP-EGFP and E) M4.GFAP-EGFP. Scale bar = 100 μm. F) Transduction specificity of the AAV serotypes in MG cell represented as the fraction of all the GFP signal that co-localized with MG marker CRALBP. N=2-3 eyes with >600 GFP signals quantified (33 for M4), error bars represent SEM. **** = p<0.0001, one-way ANOVA. Comparison of the expression pattern of GFP and the MG marker CRALBP in retinae treated with G) ShH10Y.CMV-EGFP and H) ShH10Y.GFAP-EGFP. White arrows show the co-localization of GFP signal with CRALBP+ MG cells, yellow arrows show GFP signal that was detected in other retinal cells. Scale bar = 50 μm. I) Analysis of the specificity for targeting Müller cells using the *CMV* and *GFAP* promoters. N=2-3 eyes with >600 GFP signals quantified, error bars represent SEM. **** = p<0.0001, one-way ANOVA.

Our analysis determined that the ShH10Y serotype had the best performance for transducing MG cells via IVT delivery. We next tested if pairing this capsid with a ubiquitous promoter such as CMV would retain the specificity of GFP overexpression in Müller cells. Notably, the ShH10Y.CMV-EGFP AAV could effectively target MG cells upon IVT delivery (Figure 1G). However, GFP expression was also detected in CRALBP- cells, predominantly located across the inner nuclear layer (INL), inner plexiform layer (IPL) and GCL (Figure 1G, yellow arrows). In contrast, most of the GFP+ cells in retinae treated with ShH10Y.GFAP-EGFP co-localized with CRALBP (Figure 1H). Our results showed that the substitution of *GFAP* promoter to the *CMV* promoter reduced the MG targeting specificity from 95 ± 1.6% to 47.4 ± 7.9% (Figure 1I). These results suggest that employing glial-specific promoters is important for optimizing MG targeting specificity using ShH10Y AAV. Overall, the combination of *GFAP* promoter with the ShH10 or ShH10Y capsids proved to be highly effective and specific systems for the transduction of MG cells by IVT injection.

### Exploring GLAST and HES1 as alternative promoters to target MG cells in rats

Next, we investigated two other glia specific promoters, *GLAST* ^21^ and *HES1*^22^, which could be coupled to the ShH10Y serotype to specifically target MG cells. To facilitate clinical translation, we tested the human sequence of *GLAST* and *HES1* promoters. IVT injections of ShH10Y AAV carrying different promoters were performed on SD rats to analyze AAV transduction specificity in MG cells. Our results showed that *CMV* and *GLAST* promoters drove strong GFP expression in MG cells with classical morphology (Figure 2A-B). In comparison, the *HES1* promoter also labelled some MG cells, albeit with much lower efficiency (Figure 2C). Additionally, the *CMV* promoter drove strong GFP expression in cells within the GCL and IPL layers (Figure 2A). In contrast, the *GLAST* promoter effectively drove GFP expression in CRALBP+ MG cells (Figure 2B,D). Nevertheless, GFP was also detected in CRALBP- cells within the GCL and IPL layers (Figure 2D, yellow arrows). The *HES1* promoter drove GFP expression only in a small population of retinal cells, with some CRALBP+ MG cells as well as CRALBP- cells between the GCL and the INL. (Figure 2E).

**Figure 2:**
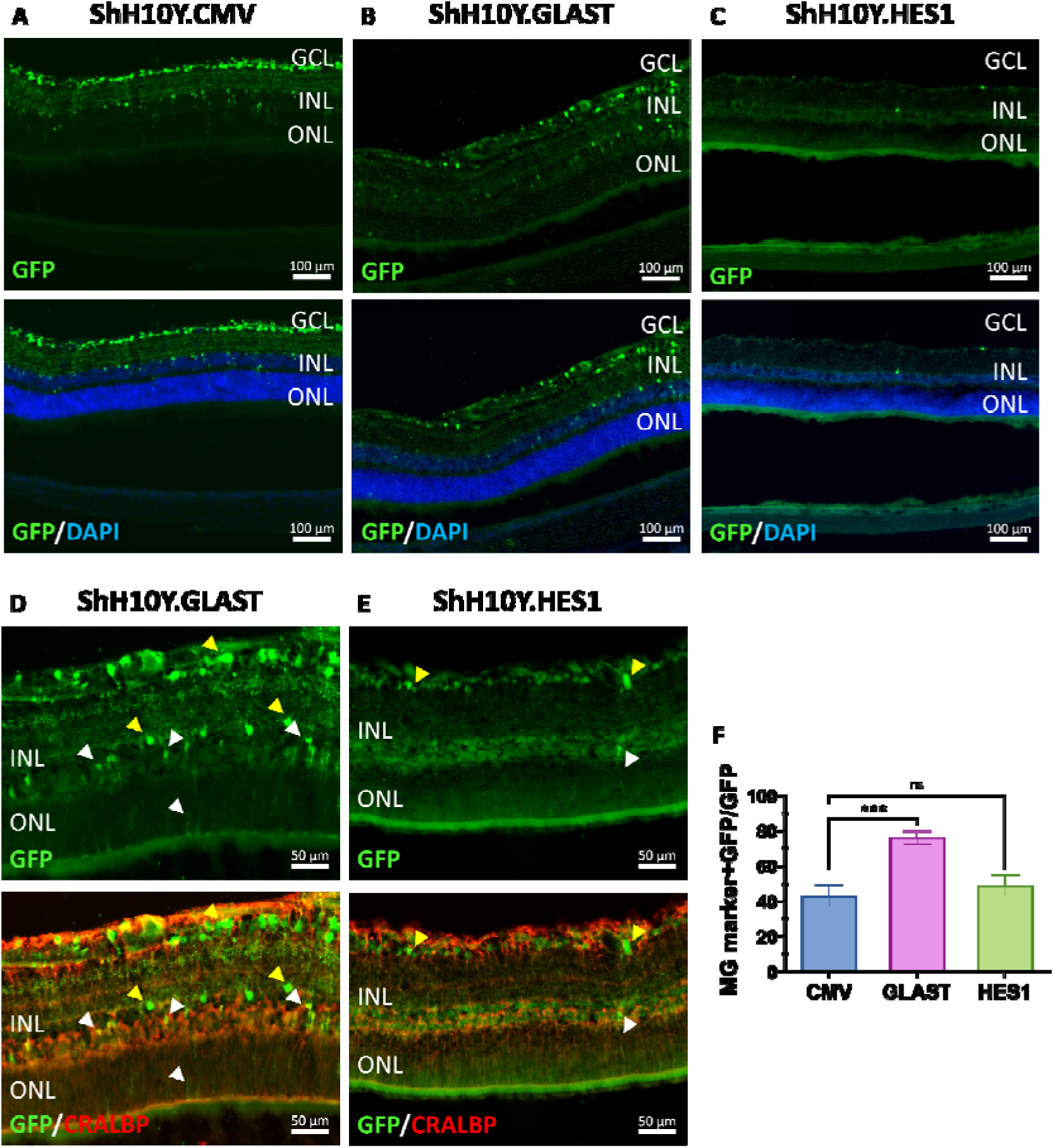
Assessment of human *GLAST* and *HES1* promoters to target MG cells *in vivo*. Representative images showing the expression pattern of GFP in rat retinae treated with IVT injections of A) ShH10Y.CMV-EGFP, B) ShH10Y.GLAST-EGFP and C) ShH10Y.HES1-EGFP. Scale bar = 100 μm. Immunohistochemistry analysis of CRALBP and GFP expression to assess promoter specificity for D) *GLAST* and E) *HES1*. White arrows show the co-localization of GFP signal with CRALBP, yellow arrows show GFP signal that was detected in other retinal cells. Scale bar = 50 μm. F) Promoter specificity analysis for *CMV*, *GLAST* and *HES1* promoters. N=2-3 eyes with >200 GFP signals quantified, error bars represent SEM. *** = p<0.001, ns = p>0.05, one-way ANOVA.

Quantification analysis confirmed that the *GLAST* promoter was significantly more specific in targeting MG cells compared to the ubiquitous promoter *CMV* (Figure 2F), with a 76 ± 3.8% specificity for *GLAST* promoter compared to 47.4 ± 7.9% specificity for *CMV* promoter. In comparison, the *HES1* promoter exhibited a specificity of 49.1 ± 5.8% in targeting MG cells. These results showed that the human *GLAST* promoter can be coupled with ShH10Y to specifically target rat MG cells, providing an interesting alternative to the *GFAP* promoter for the development of new AAV tools for gene therapy.

### Development of short GLAST and HES1 promoters to improve AAV packaging capacity

As the size of the human *GLAST* promoter we tested is large (∼2kb), we engineered shorter forms of *GLAST* promoter to improve AAV cargo capacity. Promoter composition such as enhancers, transcription factor binding sites and other regulatory elements are key factors that dictate promoter activity and tissue specificity. We performed *in silico* analysis and identified five candidate regulatory regions for the *GLAST* promoter (marked as A-E, Figure 3A). These sequences were assembled in different configurations to generate five short *GLAST* promoters: *GLAST*(ABCD) (1070 bp), *GLAST*(ABC) (839 bp), *GLAST*(BCD) (769 bp), *GLAST*(ECD) (786 bp) and *GLAST*(EABCD) (1268 bp), which were subsequently tested using the AAV expression vectors (Figure 3B, Supplementary Figure 3).

**Figure 3:**
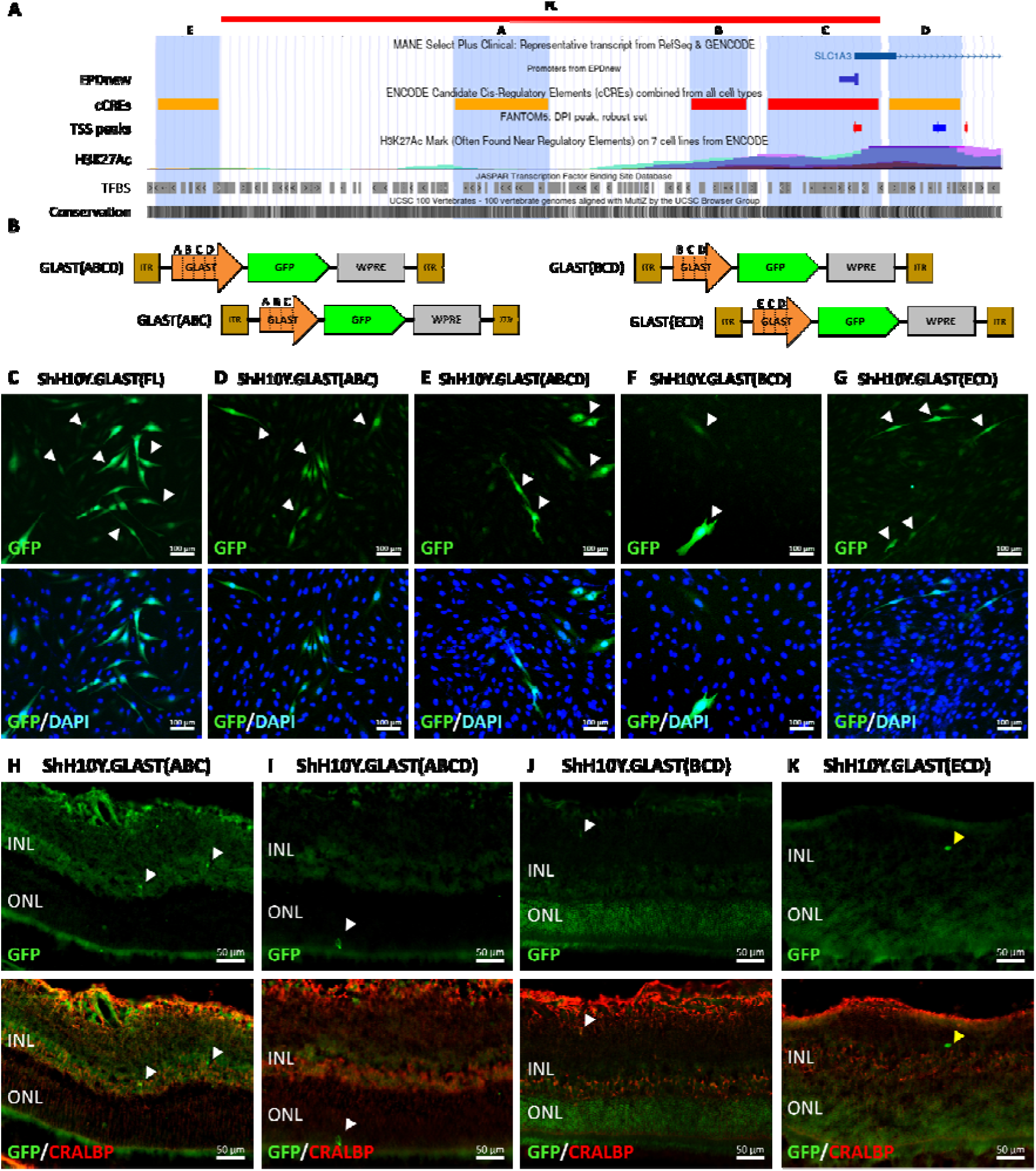
Development of short *GLAST* promoters to target MG cells. A-B) Schematic representation of the short *GLAST* promoter design. A) Five short regulatory regions from the *GLAST* promoter region (highlighted in blue, labelled A-E) were assembled to form B) the short *GLAST* promoters. FL indicated the full-length promoter. C-G) *In vitro* validation of the short *GLAST* promoters using ShH10Y AAVs to transduce the human MG cell line MIO-M1. Scale bar = 100 μm. H- K) Representative images showing the expression pattern of GFP and CRALBP in rat retinae injected with ShH10Y AAVs carrying the short *GLAST* promoter expression cassettes. White arrows show the co-localization of GFP with CRALBP and yellow arrows indicate GFP expression in other retinal cells. Scale bar = 50 μm.

We employed the human MG cell line MIO-M1 to test the functionality of these short *GLAST* promoters *in vitro*. Notably, transfection of the AAV plasmids showed that all five short *GLAST* promoters could drive GFP expression in human MG cells, with the *GLAST*(EABCD) promoter being the largest and least efficient, thus it was excluded from further studies (Supplementary Figure 3). We validated these results by packaging these vectors into ShH10Y.AAVs. Our results confirmed that ShH10Y.AAV delivery of all four short promoters could drive strong GFP expression in MIO-M1 cells, with *GLAST*(ABC) and *GLAST*(ABCD) promoters being the most efficient (Figure 3C-G).

Next, AAVs carrying the short promoters *GLAST*(ABC), *GLAST*(ABCD), *GLAST*(BCD) and *GLAST*(EBD) were delivered to rat retinae through IVT injections. Histological analysis showed that the transduction efficiency of all the AAVs injected was low, with only a few GFP+ cells per tissue section analyzed (Figure 3H-K). Specificity analysis using CRALBP expression confirmed that *GLAST*(ABC), *GLAST*(ABCD) and *GLAST*(BCD) could label MG cells *in vivo* (Figure 3H-J), but the number of GFP+ cells was too low for quantification. Also, we could not detect GFP expression in CRALBP+ MG cells using the GLAST(EBC) promoter *in vivo* (Figure 3K).

Using a similar design strategy, we constructed vectors with two short human *HES1* promoters: *HES1*(AB) of 626 bp and *HES1*(C) of 703 bp (Figure 4A-B). *In vitro* analysis of promoter activity by plasmid transfection showed that both short *HES1* promoters could drive GFP expression in MIO-M1 cells, albeit in a small number of GFP+ cells (Supplementary Figure 4A-C). This low efficiency was also observed using the full-length *HES1* promoter (Supplementary Figure 4A). We also validated the full-length and short *HES1* promoters using ShH10Y AAVs *in vitro* and obtained similar results. We detected a small number of GFP+ cells using full-length *HES1* and *HES1(AB)* promoters, while no GFP+ cells were detected using the *HES1(C)* promoter indicating the low efficiency (Figure 4 C-E). Consistent with our *in vitro* results, *in vivo* testing of ShH10Y AAVs with the short *HES1* promoters showed a limited number of GFP+ cells in the rat retina. The *HES1*(C) promoter was able to drive GFP expression in a small number of CRALBP+ MG cells, but we could not detect GFP in CRALBP+ MG cells using the *HES1*(AB) promoter (Figure 4 F-G). These results suggest that the promoter sequence of *HES1*(C) can label rat and human MG cells, but the low efficiency needs to be optimized.

**Figure 4:**
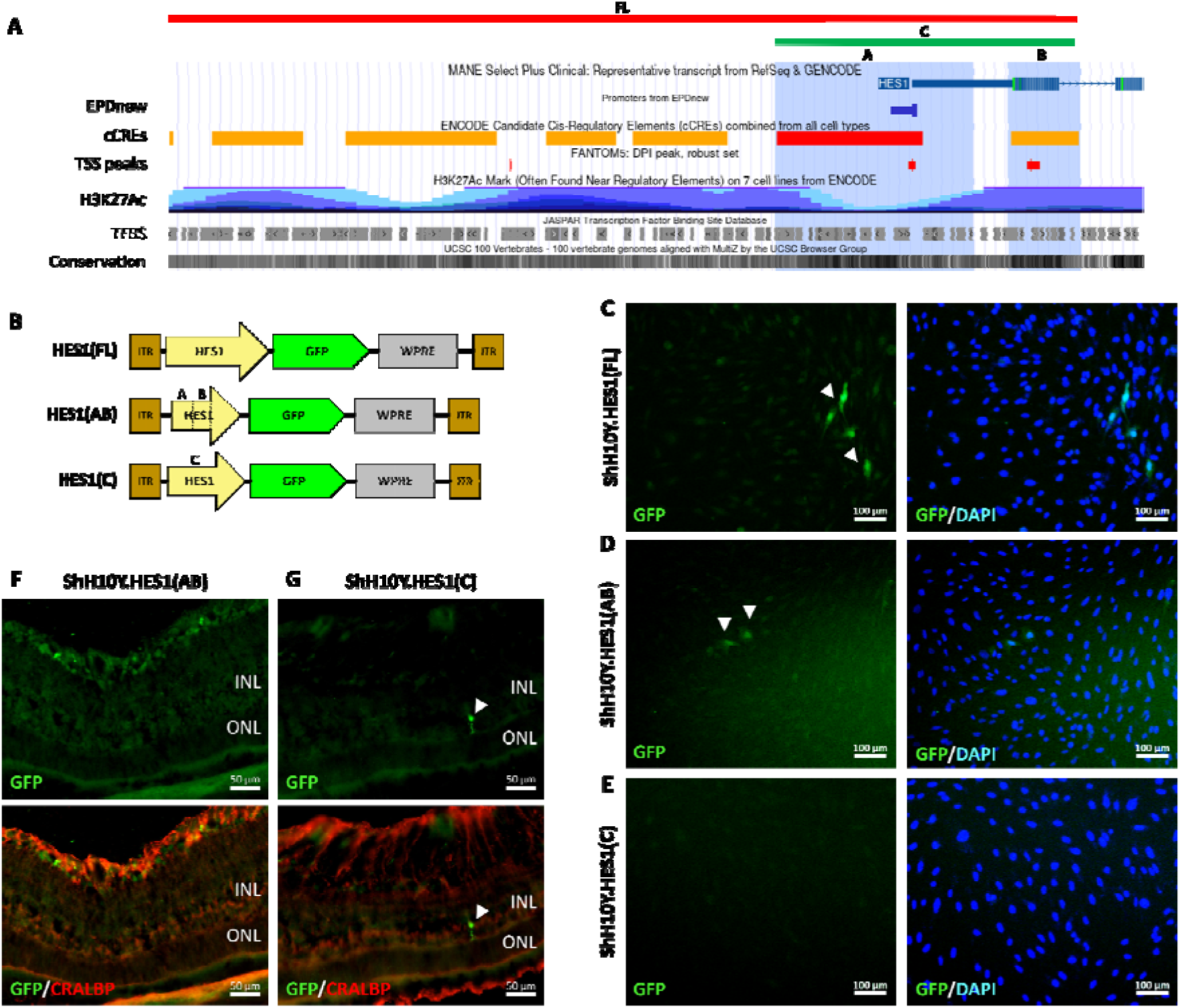
Development of short *HES1* promoters to target MG cells. A-B) Schematic representation of the human *HES1* short promoter design. A) Three putative regulatory regions from the *HES1* promoter were selected to form B) short *HES1* promoters. FL represents the full-length promoter. C-E) Validation of the short HES1 promoters *in vitro* using ShH10Y AAVs to transduce MIO-M1 cells. Scale bar = 100 μm. F-G) Representative images showing the expression pattern of GFP and CRALBP in rat retinae treated with ShH10Y AAVs carrying the short *HES1* promoter expression cassettes. White arrows show the co-localization of GFP signal with CRALBP. Scale bar = 50 μm.

Altogether, these results suggest that the short *GLAST* promoters, particularly *GLAST*(ABC), *GLAST*(ABCD), *GLAST*(BCD), can be used to target human MG cells *in vitro*, but are mostly inefficient for targeting rat MG cells *in vivo* by IVT injection. Similarly, the short *HES1*(AB) promoter showed limited activity in human MG cells *in vitro*, while the *HES1*(C) promoter showed limited efficiency for gene delivery in rat MG cells *in vivo*. These short regulatory sequences represent interesting candidates that could be further developed into valuable gene therapy tools to target MG cells.

### Identification of novel candidates for Müller glia-specific promoters

Beyond the known MG-specific promoters analyzed in this study, we used a previous single-cell RNA sequencing dataset of the human retina^23^ to identify novel promoter candidates that could specifically drive gene expression in MG cells. Differential gene expression analysis identified two novel genes that were expressed with high specificity in MG cells within the human retina: *TRDN* which encodes for Triadin, a protein involved in calcium release^24^, and *CP* which encodes for a ferroxidase that is involved in iron oxidation^25^ (Figure 5A-B). We cloned the human *TRDN* and *CP* promoter regions into our AAV-EGFP expression cassette for further testing (1.4kb and 1.9kb upstream of transcription start site respectively). *In vitro* analysis of promoter activity using MIO-M1 cells confirmed that both *TRDN* and *CP* promoters could drive GFP expression in human MG cells following plasmid transfection (Supplementary Figure 5). These results were also validated by transduction with ShH10Y AAVs carrying the same promoters *in vitro* (Figure 5C-D).

**Figure 5:**
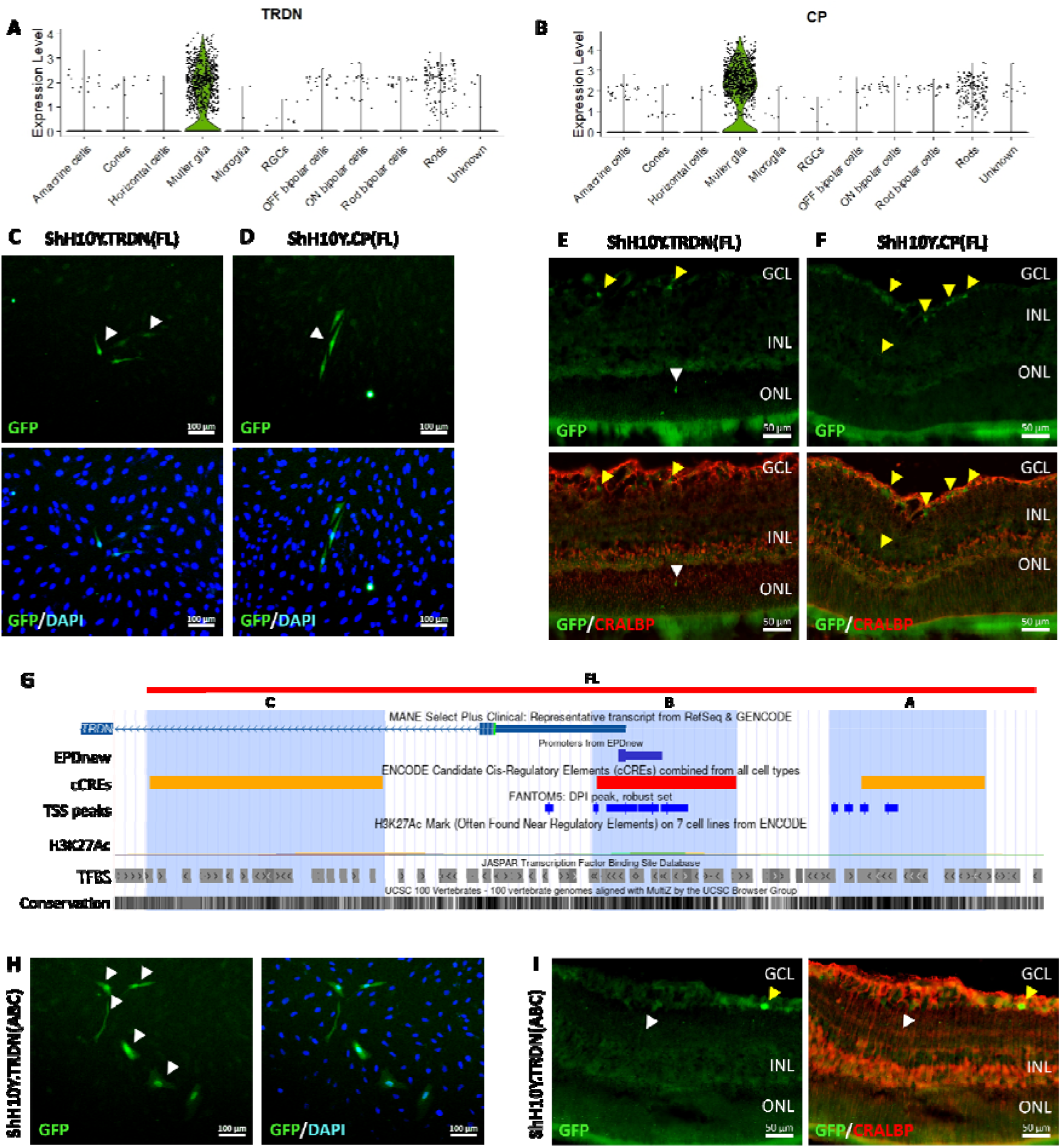
Identification and testing of novel MG-specific promoters. Differential gene expression analysis of the genes A) *TRDN* and B) *CP* in major retinal cell types in the human retina. C-D) *In vitro* validation of the *TRDN* and *CP* promoters using ShH10Y AAVs to overexpress GFP in MIO-M1 cells. Scale bar = 100 μm. Representative images showing the expression pattern of GFP and the MG marker CRALBP in rat retinae treated with AAVs carrying the E) *TRDN* and F) *CP* promoters. White arrows show the co-localization of the GFP signal with the MG marker CRALBP. Scale bar = 50 μm. G) Schematic representation of the short *TRDN* promoter design, highlighting the regulatory sequences that were assembled to form a promoter (A, B, C). H) Validation of the short promoter *TRDN*(ABC) *in vitro* using ShH10Y AAVs to target MIO-M1 cells. Scale bar = 100 μm. I) Representative images showing the expression pattern of GFP and CRALBP in rat retinae treated with ShH10Y AAVs carrying the short *TRDN* promoter expression cassette. Scale bar = 50 μm.

We performed IVT injections with ShH10Y.TRDN-EGFP and ShH10Y.CP-EGFP AAVs to test the ability of these AAV to target rat MG cells. Histological analysis showed that ShH10Y.TRDN-EGFP could label MG cells and some other retinal cells located in the GCL (Figure 5E). In contrast, AAVs carrying the CP promoter predominantly targeted cells that did not express *CRALBP* and were usually located in the GCL (Figure 5F). These results suggest that the human *TRDN* promoter has the potential to target human MG cells and to some extent for rat MG cells *in vivo*.

To optimize the *TRDN* promoter length and potentially improve efficiency and/or specificity, we designed a short version of the *TRDN* promoter. Three regulatory elements near the transcription start site of *TRDN* were assembled to form a short *TRDN*(ABC) promoter (702bp, Figure 5G). *In vitro* testing showed that the *TRDN*(ABC) promoter can drive GFP expression in MIO-M1 cells by plasmid transfection and ShH10Y AAV transduction (Figure 5H, Supplementary figure 5). Subsequent *in vivo* testing in rats showed that *TRDN*(ABC) promoter drove GFP expression in a small number of CRALBP+ MG cells (Figure 5I). These results indicate that the full-length and short *TRDN* promoters could be used for gene delivery in human MG cells *in vitro,* but further optimization is needed for *in vivo* application to target rat MG cells by IVT injections.

## Discussion

The unique properties of AAV vectors have accelerated gene therapy research, with multiple strategies harnessing the transduction profile of engineered capsids to develop treatments for inherited retinal diseases^26^. For instance, the AAV2 variant 7m8 has improved ability to penetrate and transduce all retinal layers upon IVT delivery in mouse and non-human primates (NHP)^27^, which makes this serotype an attractive tool for treatments seeking pan-retinal or photoreceptor transduction. Indeed, Ixo-vec uses 7m8 AAV to deliver the anti-VEGF drug aflibercept to the retina as a treatment for neovascular AMD^28^.

The reprogramming of MG cells into retinal neurons is a promising gene agnostic strategy to regenerate the retina and treat vision loss^29^. Thus, development of AAV tools that can specifically target MG cells is critical to realize this potential. Notably, the multilayered organisation of the retina can represent a physical barrier for AAV transduction; structures like the inner limiting membrane (ILM) and retinal neuronal processes can restrict AAV penetration^30^. ShH10 is a variant of AAV6, for which the primary target on the cell surface is sialic acid^31^. In the retina, polysialic acid can be found as a post-translational modification of the neural adhesion molecule NCAM, but it is restricted to MG cells in adult rats^32^. Also, AAV’s ability to bind heparan sulfate proteoglycans (HSPGs) influences the transduction efficiency by IVT delivery, as a diverse range of HSPGs can be found in the ILM^33^. Although AAV6 has low affinity for HSPG, capsid engineering leading to ShH10 increased affinity for HSPG and improved ability to transduce MG upon IVT injection in rodents^11^. Interestingly, Pellissier *et al*. reported some species difference in MG cell targeting specificity for ShH10 and ShH10Y viruses in mice compared to rats^20^. Our results highlighted ShH10Y as the best performing serotype to target MG cells, which is consistent with previous study^20^. Also, the selectivity of the ShH10 serotype to transduce glial cells was initially identified using human and rat tissue^11,34^, providing important tools for targeting MG cells in clinical and pre-clinical studies.

We also analyzed the transduction profile of a recently reported AAV2 variant M4 that could target human MG cells in combination with the CMV promoter^17^. This capsid showed high specificity for MG transduction in human retina explants and retinal organoids^17^. In this study, we demonstrated that the M4 serotype can transduce rat MG cells through IVT delivery with high specificity in combination with the *GFAP* promoter; however, a low transduction efficiency is observed indicating a reduced ability to transduce the rodent retina. Future studies to characterise the affinity to surface receptors and further engineering of the M4 capsid would be valuable to improve transduction efficiency across multiple species.

Our study identified the *GFAP* promoter (gfaABC1D) as the best promoter coupled with ShH10Y to target MG cells in rats *in vivo*. Previous studies in mice have used the ShH10 serotype together with *GFAP* promoter for MG reprogramming^35,36^. Interestingly, research in mice suggested that in some cases the specific transgene can alter the specificity of *GFAP* promoter, with ectopic expression observed in a portion of retinal neurons using the 7m8, PHP.eB and ShH10 serotypes^18,19^. Collectively, these reports highlight the importance of pairing MG-specific promoters and serotypes to enhance the AAV specificity to target MG cells.

In addition, we tested the specificity of other glial promoters, including *GLAST* and *HES1*. To facilitate the clinical translation of these AAV tools, we selected the human versions of these promoters for our study. The *Glast* promoter has been extensively employed to label MG cells in lineage tracing mice^18,19,37^, and the *HES1* promoter was recently used *in vitro* in reprogramming studies with human MG cells^38^. Our *in vivo* study with rat retinae further expands these findings. We showed that the IVT delivery of a combination of the *GLAST* promoter with ShH10Y was able to specifically target rat MG cells. However, the specificity achieved with this combination was still lower compared to the *GFAP* promoter. Future experiments to analyze the specificity of the human *GLAST* promoter together with ShH10Y in other clinically relevant species, such as mouse, non-human primates and human retinal explants, would allow a better understanding of the clinical value of this AAV tool for the transduction of MG cells.

In contrast, the other AAV vectors developed in this study carrying the human promoters for *HES1*, *TRDN* and *CP* were not very effective for targeting rat MG cells *in vivo*. Notably, the three promoters showed activity in human MG cells *in vitro*, though the *HES1* promoter activity was low. Of note, the *HES1* promoter was reported to be effective in human retinal explants previously^38^. This variability is possibly due to a difference in the selected *HES1* promoter sequence, as we tested a 2.5kb upstream of the transcription start site for this study. Interestingly, a 310 bp fragment located at the 3’ region of our human HES1 promoter shares 97.5% and 96.8% of homology with the mouse and rat *Hes1* promoter regions respectively, which also includes the predicted transcription start site. Future studies to characterise the functionality of the regulatory sequences in the human *HES1* promoter would provide valuable data for the generation of more AAV tools to target MG cells. Additionally, we tested two novel MG-specific promoters *CP* and *TRDN*. *CP* encodes for the protein Ceruloplasmin, which is a multicopper ferroxidase that contributes to the regulation of retinal iron and protects the cells from degeneration caused by iron overload. It has been shown that MG cells can increase the levels of *CP* expression after photopic injury^25^. On the other hand, *TRDN* is expressed in the retina (www.proteinatlas.org) and brain^39^, though its role in MG cells remains unknown. Our results showed that while the *CP* and *TRDN* promoters can drive transgene expression in human MG cells, they were not efficient in targeting the rat MG cells *in vivo* by IVT injection of ShH10Y AAVs. Further optimization of serotypes and *in vivo* delivery methods would be valuable to improve the transduction efficiency of these tools to target MG cells within the retina.

The small packaging capacity of AAVs represents a major limitation for AAV-based treatments. In particular, the reprogramming of MG cells into retinal neurons often required multiple transcription factors ^16,35^. To improve AAV packaging capacity, we engineered novel short variants of human *GLAST, HES1* and *TRDN* promoters to target MG cells. Our testing identified at least two short *GLAST* promoters that could drive gene expression in human MG cells *in vitro*. However, their ability to target rat MG cells *in vivo* is limited by IVT injection and further optimization is needed. Significant efforts have been made to develop short promoters for AAV-based retinal gene therapy^40–43^. However, many of the promoters developed for targeting MG cells are larger than 1 kb, which complicates the delivery of multiple transgenes in a single AAV. Notably, our *GLAST*(ABC) promoter is only 839 bp, which leaves enough space within the AAV vector to potentially fit multiple transgenes. Our short promoter design strategy focused on the *in silico* analysis of regulatory sequences using information from CAGE-Seq, CHIP-Seq, RNA-seq and Hi-C datasets. A similar approach was employed by Korecki *et al*. to design a short *NR2E1* promoter (1675 bp) to target mouse MG cells by IVT delivery ^40^. Future studies to test the activity, specificity of the *GLAST*(ABC) promoter across different species would be interesting, such as mouse models, NHP and human retinal explants. Overall, our results have identified potential human promoter candidates to expand the AAV toolsets to target MG cells.

This study provided a systematic testing of 4 AAV serotypes and 14 promoters to target MG cells both *in vitro* and *in vivo*. We identified the ShH10Y coupled with *GFAP* promoter as the optimal tool to target MG cells by IVT delivery, as well as other novel promoter candidates to target human MG cells, providing new resources for development of gene therapies to target MG cells.

## Methods

### Ethics

The study was approved by the Animal Ethics Committee of St Vincent’s Hospital Melbourne (024/21). Wild type Sprague Dawley (SD) rats were obtained from the Animal Resources Centre and Ozgene (Perth, Australia). The animals were housed at the Biological Research Centre (The University of Melbourne, Melbourne, Australia) or EMSU animal facility (St. Vincent’s Hospital, Melbourne, Australia).

### *Adeno-associated virus (AAV)* vector design and construction

AAV plasmids were generated through Gibson assembly using the NEBuilder HiFi DNA Assembly Master Mix (New England Biolabs, Ipswich, MA). In short, AAV expression cassette was generated with restriction enzyme digestion of the plasmid pAAV-GFAP-EGFP (a gift from Bryan Roth, Addgene # 50473). The full-length human promoters were PCR amplified from human gDNA HES1 (chr3:194,133,991-194,136,531), TRDN (chr6:123,636,303-123,637,799), CP (chr3:149,221,630- 149,224,237) and from the plasmid GLASTp-DsRed2 (a gift from Nicholas Gaiano, Addgene #17706) for GLAST (chr5:36,604,634-36,606,688), using a high-fidelity KOD Hot Start DNA Polymerase (Sigma-Aldrich, Darmstadt, Germany).

The short promoters were designed by assembling putative regulatory elements for MG-specific genes. The regulatory sequences were selected using the UCSC genome browser (https://genome-euro.ucsc.edu), taking into consideration FANTOM 5 CAGE peaks, the eukaryotic promoter database (EPDnew), transcription factor binding sites (JASPAR), open chromatin marks, CHIP-seq data and topologically associating domains (Hi-C). The short promoters were synthesized as gBlock DNA fragments (Integrated DNA Technologies, Coralville, IA) and assembled into the AAV expression cassette with EGFP reporter.

### Identification of novel MG-specific promoters

To identify MG–specific promoter usage in the human retina, we analyzed a previously reported single-cell RNA-seq dataset^25^. Single-cell RNA-seq data were analyzed using Seurat (v4.0). MG marker genes were identified using the *FindMarkers* function by comparing MG cells to all other annotated retinal cell types. Genes with log2 fold change > 0.25, pct.1 (MG cells) > 0.5, and pct.2 (all other retina cell types) < 0.05 were retained to yield high-confidence MG–specific candidates including *TRDN* and *CP*. Violin plot was used to visualize the expression of *TRDN* and *CP* across retinal cell types.

### AAV production

AAVs were generated with a triple transfection method using adherent HEK293D cells. On day 0, 2×10^7^ cells were co-transfected with 20 µg of helper plasmid (pXX6), 10 µg of RepCap plasmid and 10 µg of transfer plasmid using polyethylenimine hydrochloride (PEI; Polysciences, Warrington, PA). Three days after transfection, the AAVs were purified and concentrated using the AAVpro Purification Kit Midi (Takara Bio, Kusatsu, Japan) following the manufacturer’s instructions. The RepCap plasmids ShH10 and ShH10Y were gifts from John Flannery and Jan Wijnholds (Addgene #64867).

### In vitro testing of promoter activity

*In vitro* testing of promoter activity was performed by plasmid transfection and AAV transduction. For plasmid transfection testing, AAV plasmids carrying the specific promoter were transfected into the human MG cell line MIO-M1. In brief, the cells were maintained with DMEM high glucose supplemented with GlutaMAX and 10% fetal calf serum (Thermo Fisher Scientific, Waltham, MA). On day 0, 2.4×10^4^ cells/well were transfected with 1 µg of AAV plasmid using Lipofectamine 3000 (Thermo Fisher Scientific, Waltham, MA) following the manufacturer’s instructions. On day 1 fresh medium was added, and the samples were processed for immunocytochemistry on day 4-6. Images were taken with an Axio inverted microscope (Zeiss, Oberkochen, Germany) and processed with ZEN 3.2 Blue edition (Zeiss, Oberkochen, Germany).

For AAV transduction testing, 4×10^3^ MIO-M1 cells were transduced with AAVs with at least MOI=1,000,000 on day 0. Promoter activity was analyzed on day 6 post-transduction by immunocytochemistry to monitor GFP expression. Images were taken with an Axio inverted microscope (Zeiss, Oberkochen, Germany) and processed with ZEN 3.2 Blue edition (Zeiss, Oberkochen, Germany).

### In vivo testing of promoter activity

SD rats were anaesthetized by intraperitoneal injection of 75 mg/kg of Ketamine hydrochloride (Ceva, Hills District, Australia) and 10 mg/kg of Xylazil-20 (Ilium, Blacktown, Australia). The pupils were dilated with one drop of 1% w/v tropicamide (Bausch & Lomb, Bridgewater, NJ) and 2.5% w/v phenylephrine hydrochloride eye drops (Bausch & Lomb, Bridgewater, NJ). Intravitreal injections of AAVs were performed with a surgical microscope with the help of an intravitreal lens (OcuScience, Ann Arbor, MI) placed on the cornea. GenTeal eye lubricant gel was applied to prevent corneal desiccation. A Hamilton syringe with a 34-gauge RN needle was used to inject 3 μL of AAVs (1×10^9^ - 7×10^10^ vg) into the vitreous space of the animal eye. Two weeks post-injection, the eyes were processed for histology. The animals were euthanized with an overdose of sodium pentobarbitone (Lethabarb, Virbac, Canterbury-Bankstown, Australia). The eyes were enucleated and fixed with 4% paraformaldehyde (PFA) prior to dissection of the anterior eyecup (cornea, iris, and lens). The posterior eyecups were processed for sucrose gradient incubation (10%, 20% and 30% [w/v]) overnight. Subsequently, the tissue was embedded and snap frozen in OCT (Scigen, Paramount, CA) with an isopentane bath in liquid nitrogen. The frozen tissue samples were cryosectioned at 20 µm thickness using a CryoStar NX70 cryostat (Thermo Fisher Scientific, Waltham, MA).

### Immunohistochemistry

For immunohistochemistry analysis, the samples were permeabilized with 0.1% (v/v) TritonX-100 and blocked with 10% (w/v) normal goat serum (Sigma-Aldrich, Darmstadt, Germany). Primary antibody incubation was performed overnight at 4°C using the following dilutions: GFP (1/2000; Abcam, Cambridge, United Kingdom) and CRALBP (1/1000; Abcam, Cambridge, United Kingdom). The next day, the samples were washed with PBS and incubated with the appropriate secondary antibodies (1/400, Abcam, Cambridge, United Kingdom) for 30 min at room temperature, followed by DAPI counterstain to label the nuclei. Coverslips were mounted on the tissue sections using ProLong glass antifade mountant (Thermo Fisher Scientific, Waltham, MA). Images were taken with an Axio Imager.M2 upright microscope (Zeiss, Oberkochen, Germany) and processed with ZEN 3.2 Blue edition (Zeiss, Oberkochen, Germany).

### AAV specificity analysis

AAV specificity was analyzed by quantifying the GFP signal that co-localized with the MG marker CRALBP using the Cell Counter plugin for ImageJ. AAV specificity was calculated as a ratio of the GFP signal that co-localized with MG markers and the total GFP signal quantified. More than 10 field images corresponding to >30 sections were analyzed across 2-3 eyes treated with the AAVs, and more than 600 GFP signals were quantified per condition. The statistical analysis was performed with Graphpad Prism using one-way ANOVA and p<0.05 was used to establish statistical significance.

## Data availability

The authors confirm that data supporting the findings and conclusion of this study are presented within the article and supplemental information.

## Acknowledgements

This research was funded by the University of Melbourne (DU) and Mirugen Ltd Pty (MRFF CUREator and CUREator+). RCBW is supported by the Centre for Eye Research Australia, National Health and Medical Research Council (GNT1184076), Medical Research Future Fund (MRF2024365), Retina Australia and CASS Foundation. DU is supported by the Early Career Researcher Grant from the University of Melbourne. LW is supported by the Centre for Eye Research Australia postgraduate scholarship. The Centre for Eye Research Australia receives operational infrastructure support from the Victorian Government.

## Author contributions

DU and RCBW conceptualised and designed the study; DU, GH, LW, SC, CBN, CFL conducted experiments; DU, GH, CBN, RCBW analyzed the data; CDL, JHW, SH provided intellectual input and technical support; AWH, LL, TE, KRM, CLH, RCBW provided intellectual input and funding/material support.

## References

1. Russell, S., Bennett, J., Wellman, J.A., Chung, D.C., Yu, Z.F., Tillman, A., Wittes, J., Pappas, J., Elci, O., McCague, S., et al. (2017). Efficacy and safety of voretigene neparvovec (AAV2-hRPE65v2) in patients with RPE65-mediated inherited retinal dystrophy: a randomised, controlled, open-label, phase 3 trial. Lancet 390, 849–860.

2. McCarty, D.M., Young, S.M., Jr, and Samulski, R.J. (2004). Integration of adeno-associated virus (AAV) and recombinant AAV vectors. Annu. Rev. Genet. 38, 819–845.

3. Willett, K., and Bennett, J. (2013). Immunology of AAV-Mediated Gene Transfer in the Eye. Front. Immunol. 4, 261.

4. Issa, S.S., Shaimardanova, A.A., Solovyeva, V.V., and Rizvanov, A.A. (2023). Various AAV Serotypes and Their Applications in Gene Therapy: An Overview. Cells 12. 10.3390/cells12050785.

5. Ali, R.R., Reichel, M.B., Thrasher, A.J., Levinsky, R.J., Kinnon, C., Kanuga, N., Hunt, D.M., and Bhattacharya, S.S. (1996). Gene transfer into the mouse retina mediated by an adeno-associated viral vector. Hum. Mol. Genet. 5, 591–594.

6. Auricchio, A., Kobinger, G., Anand, V., Hildinger, M., O’Connor, E., Maguire, A.M., Wilson, J.M., and Bennett, J. (2001). Exchange of surface proteins impacts on viral vector cellular specificity and transduction characteristics: the retina as a model. Hum. Mol. Genet. 10, 3075–3081.

7. Hellström, M., Ruitenberg, M.J., Pollett, M.A., Ehlert, E.M.E., Twisk, J., Verhaagen, J., and Harvey, A.R. (2009). Cellular tropism and transduction properties of seven adeno-associated viral vector serotypes in adult retina after intravitreal injection. Gene Ther. 16, 521–532.

8. Han, I.C., Cheng, J.L., Burnight, E.R., Ralston, C.L., Fick, J.L., Thomsen, G.J., Tovar, E.F., Russell, S.R., Sohn, E.H., Mullins, R.F., et al. (2020). Retinal tropism and transduction of adeno-associated virus varies by serotype and route of delivery (intravitreal, subretinal, or suprachoroidal) in rats. Hum. Gene Ther. 31, 1288–1299.

9. Dalkara, D., Byrne, L.C., Klimczak, R.R., Visel, M., Yin, L., Merigan, W.H., Flannery, J.G., and Schaffer, D.V. (2013). In vivo-directed evolution of a new adeno-associated virus for therapeutic outer retinal gene delivery from the vitreous. Sci. Transl. Med. 5, 189ra76.

10. Pavlou, M., Schon, C., Occelli, L.M., Rossi, A., Meumann, N., Boyd, R.F., Bartoe, J.T., Siedlecki, J., Gerhardt, M.J., Babutzka, S., et al. (2021). Novel AAV capsids for intravitreal gene therapy of photoreceptor disorders. EMBO Mol. Med. 13, e13392.

11. Klimczak, R.R., Koerber, J.T., Dalkara, D., Flannery, J.G., and Schaffer, D.V. (2009). A novel adeno-associated viral variant for efficient and selective intravitreal transduction of rat Muller cells. PLoS One 4, e7467.

12. Zinn, E., Pacouret, S., Khaychuk, V., Turunen, H.T., Carvalho, L.S., Andres-Mateos, E., Shah, S., Shelke, R., Maurer, A.C., Plovie, E., et al. (2015). In Silico Reconstruction of the Viral Evolutionary Lineage Yields a Potent Gene Therapy Vector. Cell Rep. 12, 1056–1068.

13. Sherpa, T., Fimbel, S.M., Mallory, D.E., Maaswinkel, H., Spritzer, S.D., Sand, J.A., Li, L., Hyde, D.R., and Stenkamp, D.L. (2008). Ganglion cell regeneration following whole-retina destruction in zebrafish. Dev. Neurobiol. 68, 166–181.

14. Garcia-Garcia, D., Locker, M., and Perron, M. (2020). Update on Muller glia regenerative potential for retinal repair. Curr. Opin. Genet. Dev. 64, 52–59.

15. Jorstad, N.L., Wilken, M.S., Grimes, W.N., Wohl, S.G., VandenBosch, L.S., Yoshimatsu, T., Wong, R.O., Rieke, F., and Reh, T.A. (2017). Stimulation of functional neuronal regeneration from Muller glia in adult mice. Nature 548, 103–107.

16. Todd, L., Hooper, M.J., Haugan, A.K., Finkbeiner, C., Jorstad, N., Radulovich, N., Wong, C.K., Donaldson, P.C., Jenkins, W., Chen, Q., et al. (2021). Efficient stimulation of retinal regeneration from Muller glia in adult mice using combinations of proneural bHLH transcription factors. Cell Rep. 37, 109857.

17. Westhaus, A., Eamegdool, S.S., Fernando, M., Fuller-Carter, P., Brunet, A.A., Miller, A.L., Rashwan, R., Knight, M., Daniszewski, M., Lidgerwood, G.E., et al. (2023). AAV capsid bioengineering in primary human retina models. Sci. Rep. 13, 21946.

18. Le, N., Appel, H., Pannullo, N., Hoang, T., and Blackshaw, S. (2022). Ectopic insert-dependent neuronal expression of GFAP promoter-driven AAV constructs in adult mouse retina. Front. Cell Dev. Biol. 10, 914386.

19. Xie, Y., Zhou, J., Wang, L.L., Zhang, C.L., and Chen, B. (2023). New AAV tools fail to detect Neurod1-mediated neuronal conversion of Muller glia and astrocytes in vivo. EBioMedicine 90, 104531.

20. Pellissier, L.P., Hoek, R.M., Vos, R.M., Aartsen, W.M., Klimczak, R.R., Hoyng, S.A., Flannery, J.G., and Wijnholds, J. (2014). Specific tools for targeting and expression in Muller glial cells. Mol. Ther. Methods Clin. Dev. 1, 14009.

21. 21. de Melo, J., Miki, K., Rattner, A., Smallwood, P., Zibetti, C., Hirokawa, K., Monuki, E.S., Campochiaro, P.A., and Blackshawp, S. (2012). Injury-independent induction of reactive gliosis in retina by loss of function of the LIM homeodomain transcription factor Lhx2. Proc. Natl. Acad. Sci. U. S. A. 109, 4657–4662.

22. Ueno, K., Iwagawa, T., Ochiai, G., Koso, H., Nakauchi, H., Nagasaki, M., Suzuki, Y., and Watanabe, S. (2017). Analysis of Muller glia specific genes and their histone modification using Hes1-promoter driven EGFP expressing mouse. Sci. Rep. 7, 3578.

23. Gautam, P., Hamashima, K., Chen, Y., Zeng, Y., Makovoz, B., Parikh, B.H., Lee, H.Y., Lau, K.A., Su, X., Wong, R.C.B., et al. (2021). Multi-species single-cell transcriptomic analysis of ocular compartment regulons. Nat. Commun. 12, 5675.

24. Wang, Y., Li, X., Duan, H., Fulton, T.R., Eu, J.P., and Meissner, G. (2009). Altered stored calcium release in skeletal myotubes deficient of triadin and junctin. Cell Calcium 45, 29–37.

25. Chen, L., Dentchev, T., Wong, R., Hahn, P., Wen, R., Bennett, J., and Dunaief, J.L. (2003). Increased expression of ceruloplasmin in the retina following photic injury. Mol. Vis. 9, 151–158.

26. Wang, J.H., Zhan, W., Gallagher, T.L., and Gao, G. (2024). Recombinant adeno-associated virus as a delivery platform for ocular gene therapy: A comprehensive review. Mol. Ther. 32, 4185–4207.

27. Gurtsieva, D., Minskaia, E., Zhuravleva, S., Subcheva, E., Sakhibgaraeva, E., Brovin, A., Tumaev, A., and Karabelsky, A. (2024). Engineered AAV2.7m8 Serotype Shows Significantly Higher Transduction Efficiency of ARPE-19 and HEK293 Cell Lines Compared to AAV5, AAV8 and AAV9 Serotypes. Pharmaceutics 16. 10.3390/pharmaceutics16010138.

28. Khanani, A.M., Boyer, D.S., Wykoff, C.C., Regillo, C.D., Busbee, B.G., Pieramici, D., Danzig, C.J., Joondeph, B.C., Major, J.C., Jr, Turpcu, A., et al. (2024). Safety and efficacy of ixoberogene soroparvovec in neovascular age-related macular degeneration in the United States (OPTIC): a prospective, two-year, multicentre phase 1 study. EClinicalMedicine 67, 102394.

29. Woodworth, M.B., Greig, L.C., and Goldberg, J.L. (2023). Intrinsic and Induced Neuronal Regeneration in the Mammalian Retina. Antioxid. Redox Signal. 39, 1039–1052.

30. Dalkara, D., Kolstad, K.D., Caporale, N., Visel, M., Klimczak, R.R., Schaffer, D.V., and Flannery, J.G. (2009). Inner limiting membrane barriers to AAV-mediated retinal transduction from the vitreous. Mol. Ther. 17, 2096–2102.

31. Xie, Q., Lerch, T.F., Meyer, N.L., and Chapman, M.S. (2011). Structure-function analysis of receptor-binding in adeno-associated virus serotype 6 (AAV-6). Virology 420, 10–19.

32. Sawaguchi, A., Idate, Y., Ide, S., Kawano, J., Nagaike, R., Oinuma, T., and Suganuma, T. (1999). Multistratified expression of polysialic acid and its relationship to VAChT-containing neurons in the inner plexiform layer of adult rat retina. J. Histochem. Cytochem. 47, 919–928.

33. Zhang, K.Y., and Johnson, T.V. (2021). The internal limiting membrane: Roles in retinal development and implications for emerging ocular therapies. Exp. Eye Res. 206, 108545.

34. Koerber, J.T., Klimczak, R., Jang, J.H., Dalkara, D., Flannery, J.G., and Schaffer, D.V. (2009). Molecular evolution of adeno-associated virus for enhanced glial gene delivery. Mol. Ther. 17, 2088–2095.

35. Yao, K., Qiu, S., Wang, Y.V., Park, S.J.H., Mohns, E.J., Mehta, B., Liu, X., Chang, B., Zenisek, D., Crair, M.C., et al. (2018). Restoration of vision after de novo genesis of rod photoreceptors in mammalian retinas. Nature 560, 484–488.

36. Xiao, D., Jin, K., Qiu, S., Lei, Q., Huang, W., Chen, H., Su, J., Xu, Q., Xu, Z., Gou, B., et al. (2021). In vivo Regeneration of Ganglion Cells for Vision Restoration in Mammalian Retinas. Front. Cell Dev. Biol. 9, 755544.

37. Hoang, T., Kim, D.W., Appel, H., Pannullo, N.A., Leavey, P., Ozawa, M., Zheng, S., Yu, M., Peachey, N.S., and Blackshaw, S. (2022). Genetic loss of function of Ptbp1 does not induce glia-to-neuron conversion in retina. Cell Rep. 39, 110849.

38. Wohlschlegel, J., Kierney, F., Arakelian, K.L., Luxardi, G., Suvarnpradip, N., Hoffer, D., Rieke, F., Moshiri, A., and Reh, T.A. (2025). Stimulating the regenerative capacity of the human retina with proneural transcription factors in 3D cultures. Proc. Natl. Acad. Sci. U. S. A. 122, e2417228122.

39. Seo, M.H., Lim, S., and Yeo, S. (2021). Association of decreased triadin expression level with apoptosis of dopaminergic cells in Parkinson’s disease mouse model. BMC Neurosci. 22, 65.

40. Korecki, A.J., Cueva-Vargas, J.L., Fornes, O., Agostinone, J., Farkas, R.A., Hickmott, J.W., Lam, S.L., Mathelier, A., Zhou, M., Wasserman, W.W., et al. (2021). Human MiniPromoters for ocular-rAAV expression in ON bipolar, cone, corneal, endothelial, Muller glial, and PAX6 cells. Gene Ther. 28, 351–372.

41. Juttner, J., Szabo, A., Gross-Scherf, B., Morikawa, R.K., Rompani, S.B., Hantz, P., Szikra, T., Esposti, F., Cowan, C.S., Bharioke, A., et al. (2019). Targeting neuronal and glial cell types with synthetic promoter AAVs in mice, non-human primates and humans. Nat. Neurosci. 22, 1345–1356.

42. Hanlon, K.S., Chadderton, N., Palfi, A., Blanco Fernandez, A., Humphries, P., Kenna, P.F., Millington-Ward, S., and Farrar, G.J. (2017). A Novel Retinal Ganglion Cell Promoter for Utility in AAV Vectors. Front. Neurosci. 11, 521.

43. Lu, Q., Ganjawala, T.H., Ivanova, E., Cheng, J.G., Troilo, D., and Pan, Z.H. (2016). AAV-mediated transduction and targeting of retinal bipolar cells with improved mGluR6 promoters in rodents and primates. Gene Ther. 23, 680–689.

